# Genetic diversity, population structure and selection signature in Ethiopian Sorghum (*Sorghum bicolor* L. [Moench]) germplasm

**DOI:** 10.1101/2021.01.19.427274

**Authors:** Zeleke Wondimu, Hongxu Dong, Andrew H. Paterson, Walelign Worku, Kassahun Bantte

## Abstract

Ethiopia, the probable center of origin and diversity for sorghum (*Sorghum bicolor* L. [Moench]) and with unique eco-geographic features, possesses a large number of sorghum landraces that have not been well studied. Increased knowledge of this diverse germplasm through large-scale genomic characterization may contribute for understanding of evolutionary biology, and adequate use of these valuable resources from the center of origin. In this study, we characterized genetic diversity, population structure and selection signature in 304 sorghum accessions collected from diverse sorghum growing regions of Ethiopia using genotyping-by-sequencing (GBS). We identified a total of 108,107 high-quality single nucleotide polymorphism (SNPs) markers that were evenly distributed across the sorghum genome. The average gene diversity among accessions was high (H_e_ = 0.29). We detected a relatively low frequency of rare alleles (26%), highlighting the potential of this germplasm for subsequent allele mining studies through genome wide association studies (GWAS). While we found no evidence of genetic differentiation among administrative regions (F_ST_ = 0.02, *p* = 0.12), population structure and cluster analyses showed clear differentiation among six Ethiopian sorghum populations (F_ST_ = 0.28, *p* = 0.01) adapting to different environments. Analysis of SNP differentiation between the identified genetic groups revealed a total of 40 genomic regions carrying signatures of selection. These regions harbored candidate genes potentially involved in a variety of biological processes, including abiotic stress tolerance, pathogen defense and reproduction. Overall, a high level of untapped diversity for sorghum improvement remains available in Ethiopia, with patterns of diversity consistent with divergent selection on a range of adaptive characteristics.

## INTRODUCTION

Sorghum (*Sorghum bicolor* L. [Moench]), native to the dry regions of northeast Africa (Dahlberg and Wasylikowa 1996), is a major food crop in the arid and semi-arid regions of the world (Balota *et al.* 2008). It is a highly diverse crop that has experienced multiple domestication processes, resulting in five major races differentiated by inflorescence type (Harlan and Dewet 1972). Ethiopia, one of Vavilov’s centers of origin for several crop species (Vavilov 1951), hosts wide genetic variability for sorghum; all races of sorghum and their corresponding intermediates are cultivated across the country’s diverse agro-ecological zones and farming systems (Doggett 1988; Teshome *et al.* 1997; Ayana and Bekele 1998).

The wealth of genetic variability in the Ethiopian sorghum germplasm has already been noted worldwide as sources of desirable genes for sorghum improvement (Singh and Axtell 1973; Schertz 1977; Reddy *et al.* 2009). In addition, due to its unique eco-geographic features, Ethiopia possesses a large number of sorghum landraces in the gene bank as well as under subsistence agriculture. These landraces have evolved by the interaction between adaptation to a wide range of environments and selection imposed by farmers for traits enhancing agricultural productivity and performance, such as high yield, and resistance to biotic and abiotic stresses. Consequently, the genome of sorghum landraces might have experienced strong selection at genes controlling traits of agronomic and adaptive importance since domestication. Therefore, assessing genetic diversity, population structure, and selection signatures is meaningful from the perspectives of improving adequate use and conservation of these valuable resources, and may provide insights into evolutionary genomics.

Previously, genetic diversity of Ethiopian sorghum germplasm was studied using agro-morphological traits (Gebeyehu 1993; Teshome *et al.* 1997; Ayana and Bekele 1998; Ayana and Bekele 2000; Desmae *et al.* 2016b). However, this approach may not give reliable estimates of genetic diversity as these traits are limited in number and subjected to strong environmental influences (van Beuningen and Busch 1997). Genetic diversity analyses have also been carried out using various DNA marker techniques such as random amplified polymorphic DNA (RAPD) (Ayana et al. 2000), amplified fragment length polymorphisms (AFLPs) (Geleta *et al*. 2006), simple sequence repeats (SSRs) (Cuevas and Prom 2013; Adugna 2014; Desmae *et al.* 2016a; Weerasooriya *et al.* 2016), and Inter-simple sequence repeats (ISSRs) (Desmae 2007). While these studies generated useful information that is relevant to both plant breeding and germplasm conservation efforts, they were either focused on samples collected from a limited geographic range (Geleta *et al*. 2006; Desmae *et al*. 2016a), or involved limited numbers of markers (Ayana *et al.* 2000; Cuevas and Prom 2013; Adugna 2014; Weerasooriya *et al.* 2016) that are too small to fully reflect the breadth of genetic diversity that exist in the country. As a result, detailed information on genetic diversity and population structure of cultivated sorghum using reliable marker systems, while indispensible, is lacking in the center of origin, Ethiopia.

Several studies on sorghum (Hamblin *et al.* 2004; Casa *et al.* 2005; Frere *et al.* 2011; Bouchet *et al.* 2012; Mace *et al.* 2013; Morris *et al.* 2013; Zhang *et al.* 2015; Campbell *et al.* 2016; Cuevas *et al.* 2017; Tao *et al.* 2017) have utilized selective sweep analysis to detect genomic regions and genes affected by natural and artificial selection. However, most of these studies had certain limitations, either they were based on limited genome coverage (Hamblin *et al.* 2004; Casa *et al.* 2005; Frere *et al.* 2011; Bouchet *et al.* 2012) or used sorghum germplasm that have gone through the sorghum conversion program (Morris *et al.* 2013; Zhang *et al.* 2015; Cuevas *et al.* 2017). Nevertheless, these converted sorghum lines (i.e., short, early maturity and photoperiod insensitive) that are adapted to temperate regions represent partial of the genetic diversity in breeding programs, the diversity underlying traits of economic and adaptive importance remains trapped within the tropical germplasm (Cuevas and Prom 2020). Thus, characterization of the Ethiopian germplasm at genome wide scale based on patterns of nucleotide variation and selection signature will improve conservation efforts and its utilization in research and breeding programs.

Next-generation sequencing technologies have made important contributions to the development of new genotyping platforms. Genotyping-by-sequencing (GBS) is increasingly being used for profiling genome-wide nucleotide variation in many species (Elshire *et al.* 2011). The inherent characteristics of GBS including genome-wide molecular marker discovery, highly multiplexed genotyping, flexibility and low cost make it an excellent tool in genomic analysis of diverse populations, including genome-wide association studies and genomic signatures of selection (Deschamps *et al.* 2012; Poland and Rife 2012; Morris *et al.* 2013).

In this study, we used a high throughput GBS approach to generate whole genome profiles and high-quality SNP markers in a collection of 304 sorghum accessions. The objectives of this study were to (a) assess the extent and patterns of genetic diversity among sorghum accessions collected from major sorghum growing regions of Ethiopia, (b) determine the population structure of the accessions, and explore their potential for future genome wide association studies, and (c) identify genomic regions and genes potentially subjected to selection.

## MATERIALS AND METHODS

### Plant materials

A total of 304 sorghum accessions used in this study were collected from farmers’ fields of major sorghum growing administrative regions of Ethiopia (see File S1). Accessions from regions with sample size less than ten were included in adjacent regions to reduce bias due to small sample size. In doing so, six, two and two accessions from Gambella, Afar and Somalia regions were, respectively, placed under the Southern Nations, Amhara and Oromia regions. This reduced the seven regions from which the accessions were originally collected to four major sorghum producing regions (Amhara, Oromia, Tigray and South Nations). These regions include a broad swath of the range of sorghum cultivation that account for 94% of the total sorghum production in the country (Central Statistical Agency 2018). During collection readings of the coordinates and altitudes of the collection sites were recorded by a GPS map 60CSx Global Positioning System (GPS) (Garmin), which were overlaid on to the maps of Ethiopia (Figure 1).

**Figure 1.**
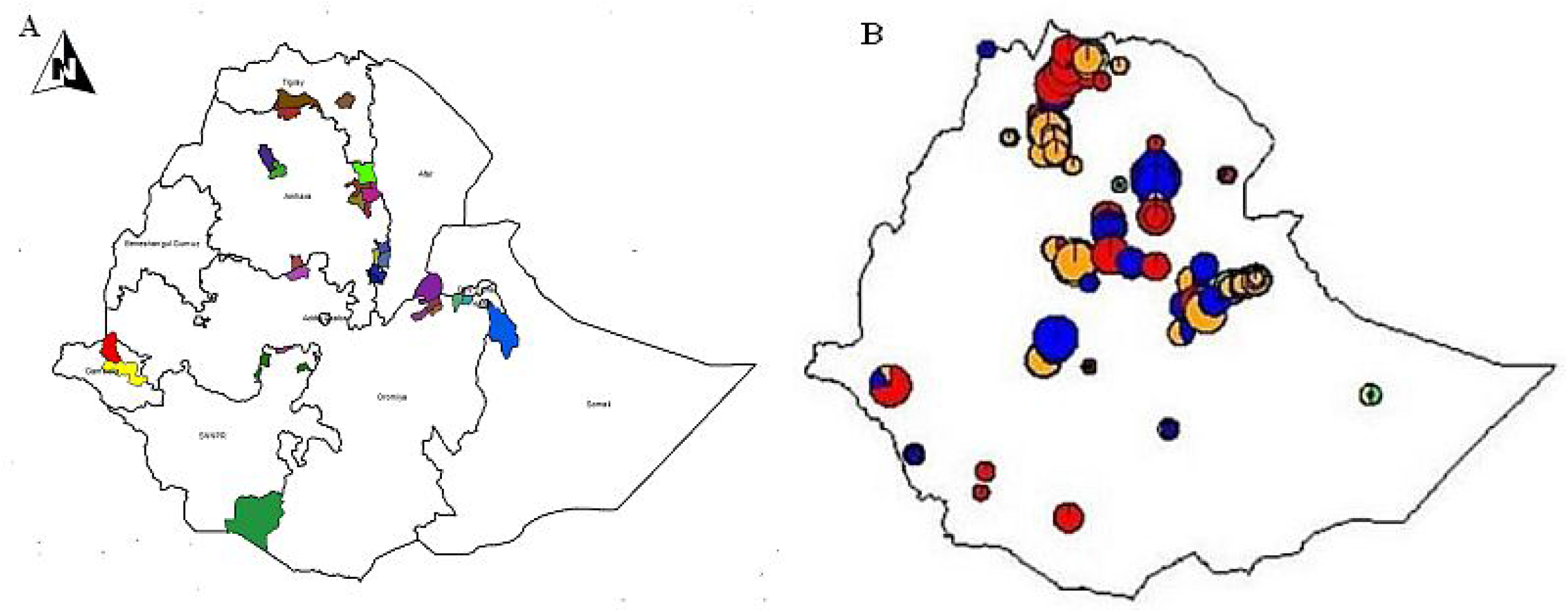
Distribution of Ethiopian sorghum accessions. (A) Geographic distribution of geo-referenced Ethiopian sorghum accessions. (B) The elevation from where each sorghum accession originated. Accessions are colored by adaptation zones (Lowland = Red; Intermediate = Blue; Highland = Orange).

### DNA extraction and genotyping-by-sequencing (GBS)

Prior to DNA extraction, the accessions were grown under field conditions and subjected to one cycle of controlled self-fertilization for purification. Leaf samples from a single representative plant per accession were collected from 15-day-old plants grown in small pots in a greenhouse. DNA was then extracted from lyophilized leaf tissues following a modified cetyltrimethyl ammonium bromide (CTAB) protocol (Mace *et al.* 2003). A total of four 96-plex GBS libraries were constructed and genotyped at the University of Georgia, Genomics and Bioinformatics Core Facility. The genotyping by sequencing (GBS) procedure (Elshire *et al.* 2011) was implemented using the *ApeKI* enzyme system. In brief, each DNA sample was digested with *ApeKI* (recognition site: G|CWCG; New England Biolabs Inc., Ipswich, MA, USA), then ligated to a unique barcoded adapter. For each library, 96 samples were pooled, and fragments with 200-500 base pair (bp) in length were extracted from a 2% agarose gel after electrophoresis and purified using a Qiagen Gel Extraction Kit (Qiagen, Hilden, Germany). The purified DNA was PCR amplified using GoTaq Colorless Master Mix (Promega, Madison, WI, USA), and the PCR product was extracted as above to eliminate primer-dimers. All libraries were sequenced on a NextSeq platform (Illumina, San Diego, CA, USA) with 150 bp single-end reads.

### Single nucleotide polymorphism (SNP) calling and quality control

SNP calling was performed using the TASSEL GBS pipeline (Bradbury *et al.* 2007) with the following parameters: kmer length of 100 bp, minimum quality score of 10, minimum call rate of 0.5, and minor allele frequency (MAF) of 0.01. Physical positions of generated SNPs were obtained based on alignment to the *Sorghum bicolor* reference genome v1.4 (Paterson *et al.* 2009). Missing data were imputed with Beagle V4.0 (Browning and Browning 2007).

### Analysis of population structure

Two approaches were used to describe the population structure of the Ethiopian sorghum collection. First, hierarchical population structure was assessed with a model-based estimation of admixed ancestry using the ADMIXTURE program (Alexander *et al.* 2009). To determine the optimal number of sub-populations (K), ADMIXTURE was run with a five-fold cross-validation (CV) procedure for K ranging from 1 to 20, and the K value with the lowest CV error was selected (Alexander *et al.* 2009). Second, pairwise genetic distances among individuals were calculated using the Sokal and Michener dissimilarity index (Sokal and Michener 1958). The resulting distance matrix was then subjected to a clustering analysis using a Neighbor-Joining (NJ) tree with 1000 bootstraps as implemented in DARwin 6.0.14 (Perrier and Jacquemoud-Collet 2006). To further investigate the spatial pattern of genetic diversity, R package tess3r was used to perform the spatial interpolation of ancestry coefficients structure onto the Ethiopian geographical map (Caye *et al.* 2016). Ancestry coefficients (*q*) estimated with ADMIXTURE program using the optimum number of subpopulations (K=6) suggested by cross-validation (CV) procedure (Alexander *et al.* 2009) were used to explore genome relatedness among 304 sorghum accessions to the stated locations of origin in Ethiopia.

### Genetic diversity and population differentiation

For each SNP, the number and frequency of alleles was calculated using TASSEL 5.0 (Bradbury *et al.* 2007). To determine the extent of genetic diversity among individuals of the entire panel, effective number of alleles (N_E_), observed heterozygosity (H_o_), gene diversity (H_e_, i.e., expected heterozygosity) and polymorphism information content (PIC) were estimated using the allele frequencies of each SNP. The above genetic diversity estimates were also computed for pooled accessions within each administrative region and ADMIXTURE inferred subpopulation. However, a comparison of diversity estimates in big populations compared that in small populations could be largely biased by the different sample sizes. To account for differences in population size, we used a subsampling scheme by taking into account the required level of precision (α = 0.05), the variances and average differences in allele frequencies between populations (Miaoulis and Michener 1976). This procedure indicated that a subsample size of 25 and 30 accessions randomly selected from each ADMIXTURE and regional population, respectively, would be appropriate to obtain unbiased estimates of the above genetic parameters (i.e., N_E_, H_o_, H_e_ and PIC) for each population. In addition, allelic richness (R_s_) and number of private alleles for each population were computed with the rarefaction method, that adjusts for differences in sample sizes across populations (Hulbert 1971), using the PopGenReport package in R (Gruber and Adamack 2014).

To estimate the components of variance among and within populations, analysis of molecular variance (AMOVA) was performed as described in Excoffier *et al.* (1992) using the R package Hierfstat (Goudet 2005). To investigate population differentiation, pairwise fixation index (F_ST_) among populations was estimated based on the method of Weir and Cockerham (Weir and Cockerham 1984) using the same package. Gene flow among populations was also estimated using indirect method based on the number of migrants per generation (N_m_) as (1-F_ST_)/4F_ST_ as described by Wright (1965).

### Detection of *F_ST_* outliers

To detect signatures of selection among ADMIXTURE subpopulations, *F_ST_* outliers were detected based on SNPs with MAF > 5% using BayeScan 2.1 (Foll and Gaggiotti 2008). To reduce the identification of false positives, a 50,000-iteration burn-in period and thinning interval size of 10 were used. The prior odd threshold to identify *F_ST_* outlier SNPs was determined using a false discovery rate (FDR) of 0.05 as implemented in the “plot_bayescan” function in R. Genes found within 100 kb of the genomic regions detected in the above test were also searched using the most recently annotated version of the sorghum genome v3.1 (www.phytozome.net). The distance 100 kb was based on the average genome-wide linkage disequilibrium (LD) decay of 100 kb (data not shown).

### Data availability

File S1 contains detailed descriptions of Ethiopian sorghum accessions, their regions of origin and geographic information. File S2 contains SNP ID numbers, locations and SNP genotypes for all accessions and SNPs. File S3 contains co-ancestry coefficient matrix of 304 Ethiopian sorghum accessions based on ADMIXTURE analysis at K = 6. File S4 contains detailed description of *F_ST_* outlier SNPs and candidate genes located in the vicinity of these SNPs based on BAYESCAN results for outlier prediction. Figure S1 contains genomic distribution of 108,107 high quality SNPs across the 10 sorghum chromosomes, and their corresponding density. Table S1 contains AMOVA and F_ST_ test results for accessions by geographic/administrative region and ADMIXTURE subgroup analyses. All supplemental materials are available at Figshare.

## RESULTS

### SNP discovery

A total of 350,618,420 reads were generated after sequencing of GBS libraries from 304 sorghum accessions. After de-duplication and alignment of unique sequence tags to the reference sorghum genome v1.4 (Paterson *et al.* 2009), a total of 236,000 SNPs were called using the GBS pipeline in TASSEL 5 (Bradbury *et al.* 2007). The quality control of SNP data (see Materials and Methods for criteria) produced a total of 115,501 high-quality SNPs (see File S2). Overall, 32.81% SNP calls were missing and imputed using Beagle V4.0 (Browning and Browning 2007). We further retained SNPs with MAF > 0.01 for downstream analysis.

A genome-wide SNP density plot (see Figure S1) revealed that the highest number of these SNPs were physically mapped on chromosome 2 (12.09%, 13,075 SNPs). The highest and lowest marker densities were observed on chromosome 7 (7.41 kb) and chromosome 5 (5.31 kb), respectively, with an average marker density of 6.13 kb per chromosome. The identified SNPs were also categorized according to nucleotide substitutions as either transitions (A↔G or C↔T) or transversions (A↔C, C↔G, A↔T, G↔T). Our analysis of transitions (Ts) and transversions (Tv) SNPs showed a Ts/Tv ratio of 1.7:1 (i.e., 68,097/40,010; Table 1), which is very close to the expected 2:1 ratio of neutral variants (Siol *et al.* 2010). The observed transition bias could be caused by a mutational bias due to intrinsic properties of DNA (e.g. cytosine deamination) in plant genomes (Gaut *et al.* 2011). Although this result suggest that most of the single nucleotide mutations observed in this study are nearly neutral, we expect that some SNPs are likely to be under selection, and thus may not susceptible to mutation bias.

**Table 1.**
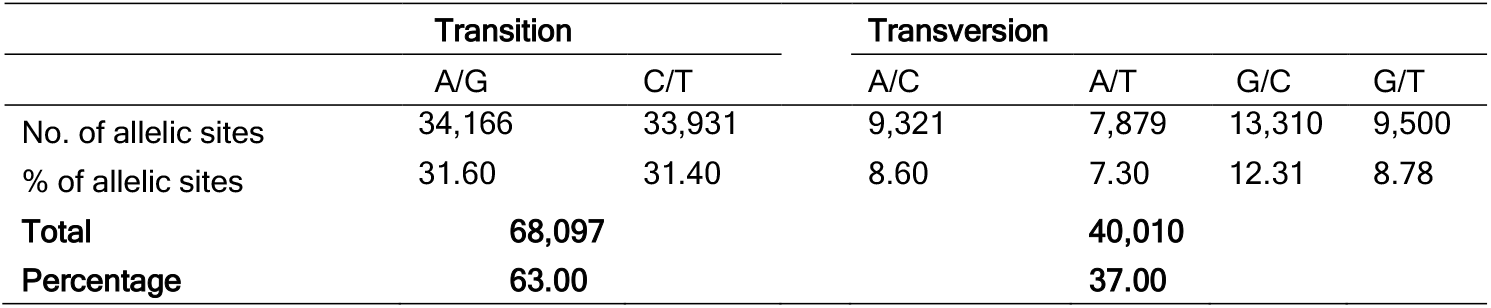
Percentage of transition and transversion SNPs identified using genotyping-by-sequencing

|  | Transition |  | Transversion |  |  |  |
| --- | --- | --- | --- | --- | --- | --- |
|  | A/G | C/T | A/C | A/T | G/C | G/T |
| No. of allelic sites | 34,166 | 33,931 | 9,321 | 7,879 | 13,310 | 9,500 |
| % of allelic sites | 31.60 | 31.40 | 8.60 | 7.30 | 12.31 | 8.78 |
| **Total** | **68,097** |  | **40,010** |  |  |  |
| **Percentage** | **63.00** |  | **37.00** |  |  |  |

### Population structure

ADMIXTURE analysis using a five-fold cross-validation (CV) for K= 1 to K = 20 indicated a steep decrease in CV error values until K = 6 (Figure 2A). For example, CV error at K = 1 was 0.57937, at K = 6 was 0.41739, at K = 8 was 0.41433, at K = 9 was 0.41027, indicating that there is no steep decrease in CV error values after K = 6. Given the modest population size in this study, we chose K = 6 as an optimal number of subpopulations, referred to as subgroups, SG-I to SG-VI (Figure 2B). Although comparing the two methods showed that there were few accessions that clustered differently depending on the analysis method, overall the clustering pattern generated using a Neighbor-Joining tree (Figure 3A), also supported the possibility that the Ethiopian sorghum collection evaluated in this study has six (K = 6) well-differentiated genetic groups and some admixtures. Therefore based on ADMIXTURE analysis, we assigned 234 (77%) accessions to one of the six subgroups with an ancestry membership coefficient probability of greater than 0.60 (q > 0.60), whereas the remaining 23% showed evidence of mixed population ancestry (see File S3).

**Figure 2.**
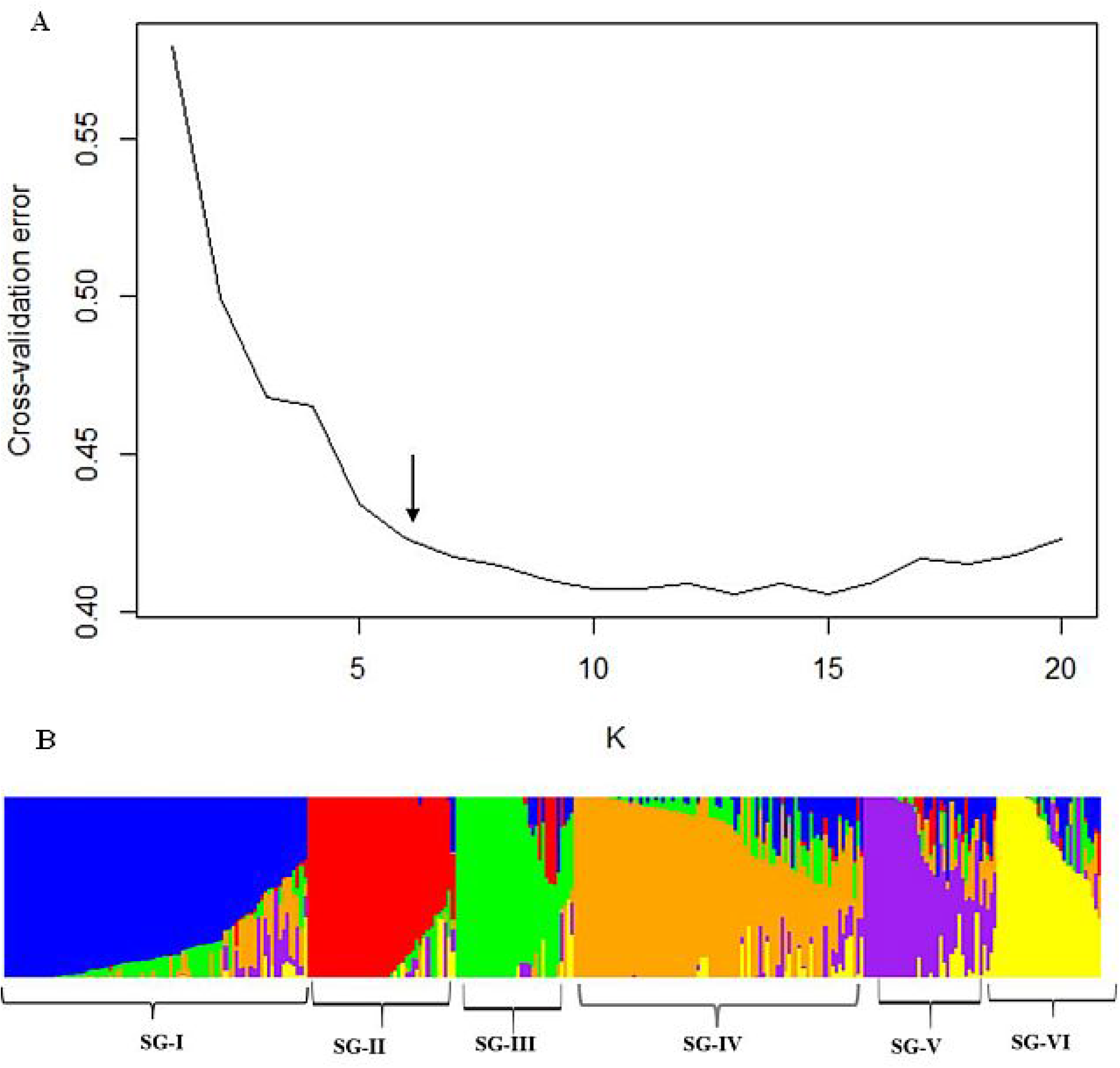
Population structure analysis of 304 Ethiopian sorghum accessions using 108,107 SNPs. (A) The cross-validation error (Y-axis) for K values from 1 to 20 (X-axis) decreased steeply until it reached 6 (arrow), suggesting an optimal number of subgroups at K = 6. (B) Bar-plot describing the population structure estimated from ADMIXTURE analysis at K = 6. Color-coding of Q-value bar plots is arbitrary. SG-I to SG-VI represents subgroup 1 to 6.

**Figure 3.**
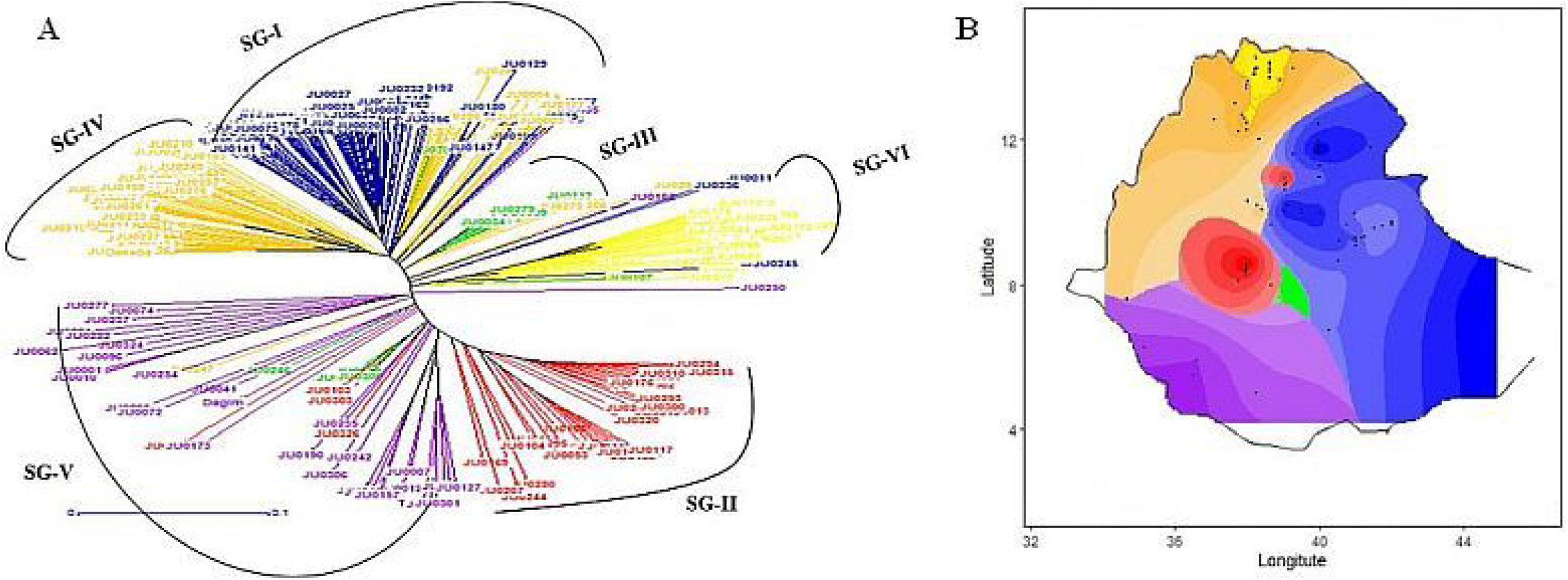
Genetic clustering of Ethiopia sorghum collection based on 108,107 SNPs. (A) Neighboring-Joining (NJ) tree of 304 sorghum accessions from DARwin 6.0.14, (B) Spatial interpolation of population ancestry coefficients across the geographic distribution of the accessions. Subgroups are color-coded based on predominant ancestry groups determined in ADMIXTURE.

According to Amede *et al.* (2015) eight agro-ecological zones (cool/humid, cool/subhumid, cool/semiarid, cool/arid, warm/humid, warm/subhumid, warm/semiarid and warm/arid) have been identified in Ethiopia based on the Global 16 Class Classification System. Given the fact that sorghum is grown in all these agro-ecologies except warm/humid and cool/arid (Menamo *et al.* 2020), we hypothesized that genetic groups could reflect to population structure across the agro-ecological zones of Ethiopia. Consistent with this hypothesis, we observed strong geographic clustering when the accessions were mapped by group (Figure 3B), suggesting significant contribution of agro-ecological variation to ancestry, with individuals in specific subgroup found to co-locate in geographic regions. For instance, in SG-I (blue), the majority of the individuals were from eastern parts of Ethiopia (Figure 3B). In addition, individuals with high membership coefficients in SG-III (green), SG-V (purple) and SG-VI (yellow) showed strong clustering according to their geographic origin (central, western and northern parts of the country, respectively). In contrast, SG-II which includes 41 accessions showed modest clustering according to geography, suggesting that the population structure could also be affected by other factors such as seed exchange and food preferences (Deu *et al.* 2010).

### Genetic diversity and population differentiation

Based on the allele frequency distribution of this collection, 26% of the SNPs were rare (MAF < 0.05) (Figure 4). For the four regional populations, we found a similar proportion of rare alleles, with values ranging from 15% to 20% in accessions collected from the Amhara and Tigray regions, respectively. Among the six ADMIXTURE subgroups, SG-III had the highest percentage (46%) of rare alleles, while in the remaining groups (SG-I, SG-II, IV, V and VI), an average of 24% of the detected alleles were rare. The distribution of these rare alleles among populations could represent a recent admixture as a result of inter-population gene flow (Memon *et al.* 2016). The correlation of a recent admixture and the distribution of rare alleles among populations, may in part explained by the fact that some alleles would have the opportunity to be reintroduced to the populations through recent introgression, and that these alleles will come to be spread fairly evenly in the populations in very small quantities. In addition, pairwise population comparisons showed that 76% to 91% of the SNPs common in at least one population were common among all regional populations (i.e., 58,673 SNPs), whereas only 18% to 41% of such SNPs were common among all ADMIXTURE subgroups (i.e., 12,400 SNPs).

**Figure 4.**
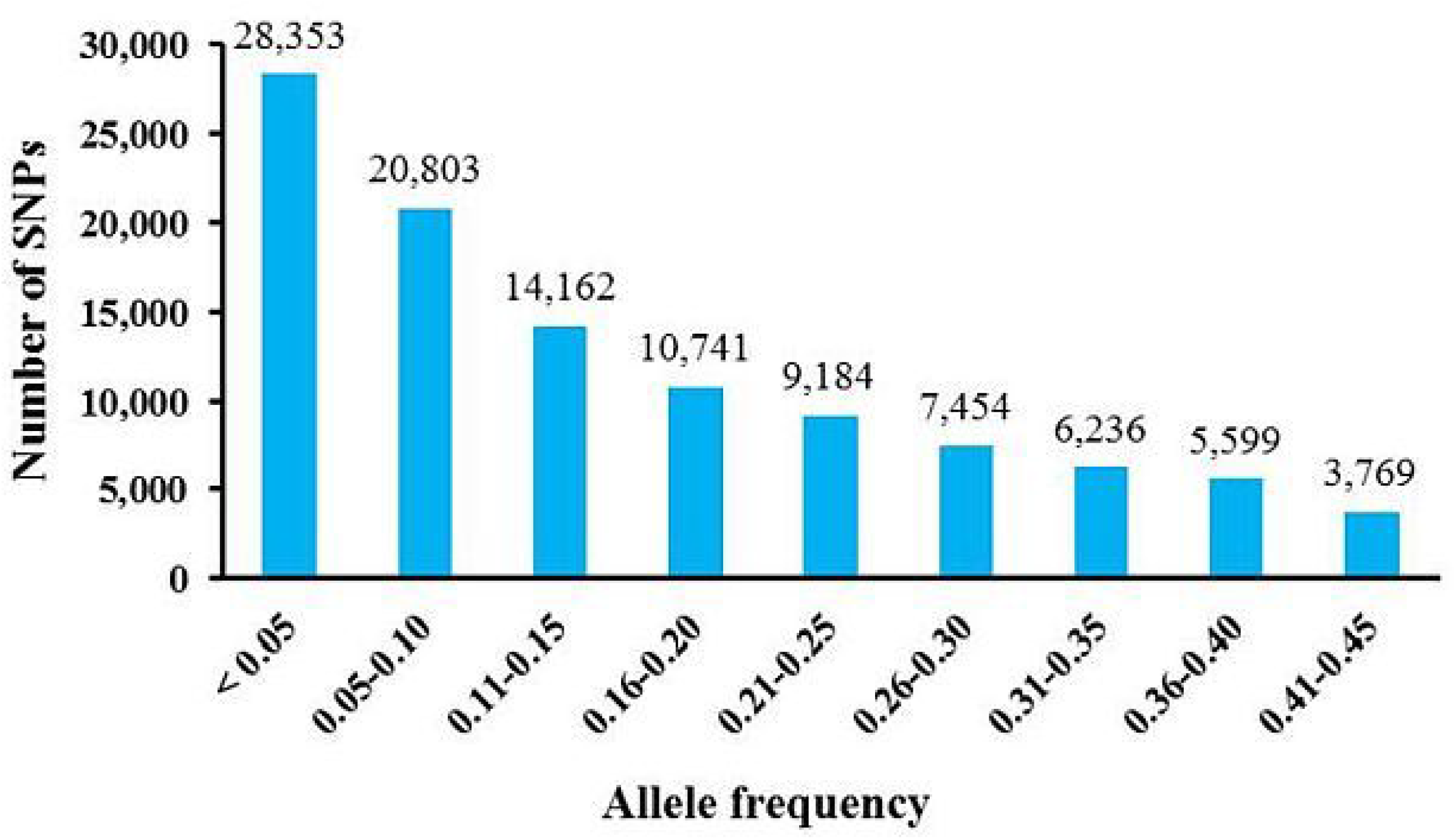
Minor allele frequency (MAF) and number of SNPs based on 304 sorghum accessions from Ethiopia.

Genetic diversity parameters for the entire panel, regional populations and ADMIXTURE subgroups are summarized in Table 2. The individual SNP PIC values of the entire panel ranged between 0.09 and 0.37, with an average value of 0.24 across all polymorphic loci (Table 2). Gene diversity (H_e_; i.e. expected heterozygosity) ranged from 0.09 to 0.50, and its value average all loci was 0.29 (Table 2). The mean observed heterozygosity value (H_o_ = 0.12) of the entire panel was similar with that observed in previous studies of sorghum landraces (Dje *et al.* 2004; Cuevas *et al.* 2017). Among the four regions, the highest level of genetic diversity was observed in accessions collected from the Tigray region (N_E_ = 1.53, H_e_ = 0.32, PIC = 0.26), and the lowest in the Oromia region (N_E_ = 1.42, H_e_ = 0.28, PIC = 0.24). Allelic richness based on rarefaction was relatively higher in the Southern Nations (R_S_ = 1.81) and lower in the Oromia region (R_S_ = 1.77) (Table 2). Considering the genetic diversity among the six ADMIXTURE subgroups, the highest level of genetic diversity (N_E_ = 1.52, R_S_ = 1.77, H_e_ = 0.32) was found in SG-V (Table 2). SG-VI had the second highest level of diversity (R_S_ = 1.66, H_e_ = 0.31) and harbored many private alleles (N_PA_ = 1,430) compared to other subgroups (Table 2).

**Table 2.** Genetic diversity estimates of Ethiopian sorghum accessions at different population levels

| Population | N<sup>a</sup> | n | N<sub>E</sub> | R<sub>S</sub> | N<sub>PA</sub> | H<sub>o</sub> | H<sub>e</sub> | PIC |
| --- | --- | --- | --- | --- | --- | --- | --- | --- |
| Entire panel | 304 | - | 1.46 | - | - | 0.12 | 0.29 | 0.24 |
| **Administrative regions** |  |  |  |  |  |  |  |  |
| Amhara | 146 | 30 | 1.47 | 1.79 | 776 | 0.13 | 0.30 | 0.24 |
| Oromia | 77 | 30 | 1.42 | 1.77 | 230 | 0.11 | 0.28 | 0.24 |
| Tigray | 44 | 30 | 1.53 | 1.78 | 126 | 0.19 | 0.32 | 0.26 |
| Southern Nations | 37 | 30 | 1.48 | 1.81 | 55 | 0.12 | 0.30 | 0.24 |
| **ADMIXTURE subgroups** |  |  |  |  |  |  |  |  |
| SG-I<sup>b</sup> | 84 | 25 | 1.51 | 1.60 | 683 | 0.19 | 0.30 | 0.26 |
| SG-II | 41 | 25 | 1.50 | 1.50 | 500 | 0.18 | 0.29 | 0.25 |
| SG-III | 33 | 25 | 1.32 | 1.33 | 12 | 0.01 | 0.23 | 0.20 |
| SG-IV | 80 | 25 | 1.51 | 1.64 | 648 | 0.19 | 0.30 | 0.26 |
| SG-V | 37 | 25 | 1.52 | 1.77 | 637 | 0.15 | 0.32 | 0.26 |
| SG-VI | 29 | 25 | 1.50 | 1.66 | 1430 | 0.15 | 0.31 | 0.25 |

<sup>a</sup>N: Number of individuals; n: Subsample size; N<sub>E</sub>: Effective number of alleles; R<sub>S</sub>: Allelic richness, N<sub>PA</sub>: Number of private alleles, H<sub>o</sub>: Observed heterozygosity, H<sub>e</sub>: Expected heterozygosity, PIC: Polymorphism information content and F<sub>IS</sub>: Inbreeding coefficient; <sup>b</sup> SG-I to SG-VI represents subgroup 1 to 6. Except for the entire panel, basic diversity statistics (N<sub>E</sub>; H<sub>o</sub>; H<sub>e</sub> and PIC) were estimated based on subsamples of equal size.

Analysis of molecular variance (AMOVA) showed that 98% of the total variation was found within regions, while 28.06% and 71.94% of the total variation was found among and within ADMIXTURE subgroups, respectively (see Table S1). The number of migrants per generation as indirect estimate of gene flow was also very high (N_m_ = 12.25) among the regions, leading to a low genetic differentiation between the regions. The pairwise fixation index (F_ST_) among ADMIXTURE subgroups ranged from 0.11 to 0.48, indicating a relatively high level of genetic differentiation (Table 3) that resulted from a restricted gene flow (N_m_ = 0.64) among the populations.

**Table 3.** Pair wise F_ST_ matrix, a measure of population divergence among the six ADMIXTURE subgroups

|  | SG-II<sup>a</sup> | SG-III | SG-IV | SG-V | SG-VI |
| --- | --- | --- | --- | --- | --- |
| SG-I | 0.44 | 0.18 | 0.11 | 0.25 | 0.31 |
| SG-II |  | 0.48 | 0.41 | 0.21 | 0.23 |
| SG-III |  |  | 0.16 | 0.27 | 0.38 |
| SG-IV |  |  |  | 0.21 | 0.29 |
| SG-V |  |  |  |  | 0.23 |

<sup>a</sup> SG-I to SG-VI represents subgroup 1 to 6

### Genomic signatures of selection

The identification of functional genomic regions that might be targets of selection provides information useful for the discovery of candidate genes of breeding importance in sorghum (Campbell *et al.* 2016). In this study, a total of 79,754 SNPs (MAF > 0.05) were tested for evidence of selection among the six ADMIXTURE subgroups using BayeScan v.2.1 (Foll and Gaggiotti 2008). This approach distinguishes between loci that diverged via random drift and those that diverged via selection. Among the 79,754 SNPs analyzed, 40 (FDR < 0.05) present evidence of selection among the six genetic groups according with BAYESCAN results (Figure 5; see File S4). Among these 40 *F_ST_* outlier SNPs, 38 were consistent with the evidence of diversifying selection (α > 0) and two corresponded to balancing selection (α < 0).

**Figure 5.**
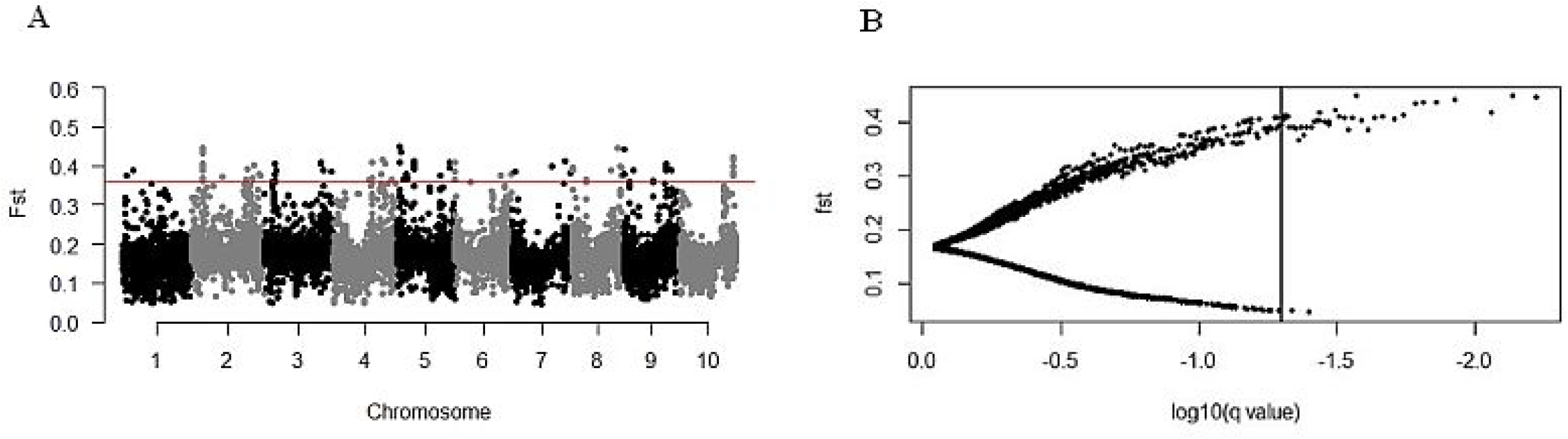
BAYESCAN results for the analysis of 79,754 SNPs among six Ethiopian sorghum subgroups for outlier prediction. (A) The distribution of *F_ST_* values across the sorghum genome. The red horizontal line is a cut-off for top *F_ST_* outlier SNPs. (B) Each *F_ST_* value is plotted against the log10 of the corresponding q-value for outlier prediction. The vertical line indicates the threshold false discovery rate (FDR = 0.05) value used to identify outlier SNPs represented on the right side of the line.

We also identified a total of 47 candidate genes in the vicinity of the genomic regions containing these *F_ST_* outlier SNPs (see File S4). These candidate genes represented different categories of biological processes, including regulation of biotic and abiotic stress tolerance (F-box proteins, MADS box transcription factor), signal transduction (similar to low temperature-responsive RNA-binding protein), plant cell wall synthesis (Glycosyl transferase 1) and ion transport.

## DISCUSSION

Ethiopia, the probable center of origin and diversity for sorghum and with unique eco-geographic features, possesses a large number of landraces in the gene bank as well as under subsistence agriculture. The germplasm from this region represents one of the most important sources of useful genes for sorghum improvement efforts around the world (Dogget 1988; Reddy *et al.* 2009; Adugna 2014). In addition to providing a broad sample of the diversity in sorghum, the genotypes included in this study are known to display agronomically important traits including drought tolerance (Wondimu *et al.* 2020). Therefore, the genomic characterization presented herein provides an advantageous starting point to make adequate use of these valuable resources, and could also be employed for the genomic dissection of important phenotypes in sorghum.

### Genetic diversity and regional differentiation

In this study, a high-throughput GBS technology was used to explore genetic diversity, population structure, and selection signature in sorghum accessions collected across the center of origin and domestication, Ethiopia. Indeed, the lower frequency of rare alleles (Figure 4) observed in our study than in a previous GBS analysis of Ethiopian sorghum landraces (Girma *et al.* 2019), highlights the potential of this collection for subsequent allele mining studies through GWAS. The level of diversity in this Ethiopian collection is higher than that observed in the global sorghum association panel (Maina *et al.* 2018), which confirms Doggett’s long standing hypothesis that Ethiopia is not only part of the center of origin but also the center of diversity of sorghum (Doggett 1988). The diverse agro-ecological zones and farming systems where sorghum is grown in Ethiopia as well as the high level of gene flow between cultivated sorghum and its wild relatives, all seem to have contributed to the wide range of variation observed in this and previous studies (Snowden 1936; Stemler *et al.* 1977; Teshome *et al.* 1997; Ayana and Bekele 1998, 1999; Tesso *et al.* 2008) of Ethiopian sorghum germplasm. An intriguing hypothesis is that the richness of diversity in Ethiopia may facilitate selection for different allele combinations that result in particular suites of traits, providing rich genetic sources for sorghum improvement programs.

Our results (see Table S1) support previous reports of low level of regional differentiation for cultivated sorghum in Ethiopia (Ayana and Bekele 1998; Ayana *et al.* 2000; Desmae *et al.* 2016a; Desmae *et al.* 2016b). The lack of regional differentiation could be attributed, at least in part, to frequent gene flow as a consequence of extensive exchange of materials between farmers from these regions, which was also confirmed by the high rate of gene flow observed among the regions. An alternative or perhaps complementary explanation for the lack of regional differentiation is that these regions do not represent different agro-environmental conditions but political regions formed based on the federal system of Ethiopia. Overall, these results suggest that a single large random collection from the whole area would be adequate to capture and preserve most of the genetic variation present in Ethiopian sorghum germplasm. However, the high level of allelic diversity in population from the Southern Nations (Table 2) could be an indicator of the conservation status of its genetic diversity, thus additional collection from this region may be needed to support and increase the genetic diversity of Ethiopian sorghum germplasm collection.

### The structure of genomic diversity in Ethiopian sorghum

While regional differentiation is lacking, analysis of genotypic data revealed clear differentiation among six Ethiopian populations (Figures 2 and 3). Previous studies have shown that sorghum populations are structured according to botanical races and geography (Barnaud *et al.* 2007; Deu *et al.* 2010; Maina *et al.* 2018). Since the accessions used in the current study were not characterized for racial groups, it was not possible to relate the observed genetic structure with racial category. Of the different sorghum growing agro-ecological regions in Ethiopia, the wetter regions mostly represent the western parts of the country which receive high rainfall, and with rainfall rapidly decreasing to the east (Amede *et al.* 2015). Our spatial analysis (Figure 3B) separates populations in the SG-V (purple; mostly from the western parts of Ethiopia) from SG-I (blue; predominantly found in the eastern parts of the country), consistent with these two populations inhabiting contrasting environments, at least in terms of rainfall. Surprisingly, we also observed distinct geographic groupings between subgroups III (green) and IV (orange). SG-III is found in central Ethiopia, which is mostly characterized by cool/subhumid conditions (Amede *et al.* 2015), while SG-IV mainly from the northeastern parts of the country that is generally characterized by warm/semiarid conditions, providing additional insights into the patterns of ancestry resulting from adaptation to different agro-ecologies. In contrast, SG-II (red) is found in multiple groups along with populations from SG-I (blue), suggesting that the population structure may also be affected by other factors such as human activities including seed exchange and food preferences (Deu *et al.* 2010).

Overall, the observation that the six distinct genetic groups identified equaled the number of different agro-ecological zones (i.e., cool/humid, cool/subhumid, cool/semiarid, warm/subhumid, warm/semiarid and warm/arid) of Ethiopian sorghum may indicate a strong contribution of agro-ecological variation to genetic groupings observed in this study. Further studies that combine population analyses with environmental and phenotypic trait variables should provide a more complete understanding of sorghum genetic structure in Ethiopia and facilitate breeding of locally adapted sorghum varieties.

### Signatures of selection

We expect higher genetic population differentiation for adaptive SNP than neutral SNP if adaptation to local environments is the principal source of genetic differentiation (Villemereuil and Gaggiotti, 2015). To identify genomic regions that may be under selection pressure, we used the *F_ST_* outlier method (see Materials and Methods section). Of the 40 outlier SNPs identified, 31 were located at less than 100 kb from annotated genes (see File S4), which provides the first support for their putative relevance.

For instance, the top *F_ST_* outlier SNP (S5_3030678; *F_ST_* = 0.45) with a signature of diversifying selection, was located at ~ 24.42 kb from Sobic.005G033801, a candidate gene which encodes flavonol 3-O-glucosyltransferase protein known to be involved in anthocyanin biosynthesis pathway (Holton and Cornish 1995). Anthocyanin accumulation appeared to be associated with grain pericarp color (Awika *et al.* 2004, 2005), and protection against bird predation (Xie *et al.* 2019) in sorghum. However, the presence of pericarp color is undesirable in sorghum grains for making *injera* (a traditional pancake like bread in Ethiopia) for human consumption (Ayana and Bekel 1998). The diversifying selection observed here thus could be the result of opposing selection pressures driven by humans and natural conditions. We found two additional candidate genes (Sobic.005G041200 and Sobic.007G135301) located at 64.13 kb and 17.33 kb from S5_3724645 and S7_56063576 on chromosome 5 and 7, respectively. The Sobic.005G041200 gene encodes F-box protein known to be associated with sorghum response to various stresses including drought (Johnson *et al.* 2014). While the Sobic.007G135301 gene encodes MADS box protein which has been associated to smut resistance in sorghum (Girma *et al.* 2019) and maize (Wang *et al.* 2012). Another candidate gene (Sobic.003G269600), for which evidence of association with sorghum phenotypic diversity for plant height had been reported (Phuong *et al.* 2013), was identified near S3_60642776. In Ethiopia, sorghum grows in diverse agro-ecologies ranging from the hot and dry lowlands to high-altitude regions (Snowden 1936), where different environmental conditions favor different biotic and abiotic stress factors. Thus, the diversifying selections detected in this study are expected, as the type of selection acting on a gene can be different between populations depending on the environments.

## CONCLUSION

This study reported a high level of genetic diversity and differentiation among Ethiopian sorghum accessions, which provides a great opportunity for developing new cultivars with desirable characteristics. Our results illustrate how populations adapting to different environments become structured genetically on small spatial scales. We found genomic regions of potential interest, with further large-scale phenotypic and geographic characterization can provide multiple lines of evidence for the putative importance of these particular loci in the genetic control of traits of economic and adaptive importance in sorghum. Overall, this study contributes to the genomic resources available for sorghum improvement efforts around the world.

## ACKNOWLEDGEMENTS

We are grateful to the USAID’s Feed the Future Laboratory for Climate Resilient Sorghum for providing financial support to undertake this study. We also thank the staff of Genomics and Bioinformatics Core Facility at the University of Georgia for their support in GBS and bioinformatics services.

